# Mobile Phone Based Strategies for Preconception Education in Rural Africa

**DOI:** 10.1101/586636

**Authors:** Zemenu Yohannes Kassa, Zelalem Tenaw, Ayalew Astatkie, Melese Siyoum, Gezahegn Bekele, Kefyalew Taye, Shewangizaw Mekonnen, Zerai Kassaye

## Abstract

**Background:** prepregnancy health care is vital to alleviate and prevent maternal and neonatal disability and death.

**Objective:** The purpose of the study was to measure the levels of knowledge and attitude on preconception care and their determinants among women who delivered at government hospitals in a rural setting in southern Ethiopia.

**Method:** A facility-based cross sectional study was done from January 01 to February 30, 2017 on a sample of 370 women who delivered at government hospitals in Wolayita zone. The mothers were selected using systematic random sampling. The data were collected using structured and pretested interviewer administered questionnaires at the postnatal ward of each hospital. Data were analyzed using bivariate and multivariable techniques.

**Results:** The result showed that 53% (95% confidence interval [CI]: 47.8%, 58.1%) of mothers who delivered at public hospitals had adequate level of knowledge on preconception care, whereas 54.3% (95% CI: 49.2%, 59.5%) possessed positive attitude to preconception care. Mothers who have radio, planned pregnancy and have participated in community meetings related to preconception care had a meaningfully higher odds of good level of knowledge to preconception care. Ordinal regression showed that women who own mobile phone had at least three times significantly higher odds of positive attitude to preconception care, whereas women who have participated community meetings had lower odds of positive attitude on preconception care.

**Conclusion:** The results revealed that the levels of mothers’ knowledge and positive attitude on preconception care are low relative to other studies. Using transistor radio and mobile phone have significant effect in improving the knowledge and attitude of reproductive age women on preconception care. Hence, providing community health education based on radio and/or mobile phone messaging could be useful in positively influencing the knowledge and attitude of women on preconception care.

## Background

Preconception health care is a set of interventions prepregnancy to reduce the influence of biomedical, behavioral and social risks of mothers’ health, and unborn child health (1). It can improve maternal and neonatal outcome by identifying, modifying bad habits and behaviors before conception and decreasing unintended pregnancies(2). Besides, most of pregnancy and childbirth complications can be alleviated by implementation of preconception care at health institution, meanwhile in low resource settings preconception care is not regularly implemented (3).

Though both governments and civil societies in developing counties frontline agenda is maternal and neonatal health service, newborn and child death and stillbirth, of which 77% are preventable by creating platform for essential packages at community, health center and hospital levels, have not yet been reduced to the expected level (4).

Worldwide 216 maternal deaths occurred per 100,000 live births in 2015, of which 99% occurred resource constraint areas, especially south Asia and sub-Saharan Africa. The most substantial cause of women mortality are: obstetric hemorrhage, preexisting medical conditions, hypertensive disease of pregnancy, infections /sepsis, unsafe abortion, and other indirect causes (4). Globally, 2.6 million children died in the first month of life and neonatal mortality was estimated at 19 deaths per 1,000 live births(5). The under-five mortality in 2015 was 42.5 per 1,000 live births (6, 7). In Ethiopia maternal mortality ratio is estimated at 412 per 100,000 live births in 2016, neonatal mortality at 29 deaths per 1000 live births, infant mortality at 48 deaths per 1,000 live births and the under-five mortality at 67 per 1000 live birth in 2016(8).

Reproductive planning through preconception care could be reduced 71% of unwanted pregnancies, thereby eliminating 22 million unplanned births, 25 million induced abortions and 7 million miscarriages(9, 10). Similarly lack of preconception care and low folic acid supplementation for women in developing countries might increase the risk of neural tube defect in newborns by four time compared with developed countries (11).

The basic concept of preconception care is to advise child bearing age women away from any negative health behaviors or conditions that might affect a future pregnancy(12). “A reproductive health plan reflects a person’s intentions regarding the number and timing of pregnancies in the context of their personal values and life goals”. This health plan will increase the number of planned pregnancies and encouraged persons to address risk behaviors before conception, reducing the risk of adverse outcomes for both the mother and unborn child(13, 14).

A study done in Kelantan, Malaysia found that 51.9% of women attending maternal health clinic had good level of knowledge on preconception care and 98.5% had positive attitude on preconception care(15). A study done in Egypt revealed that 39.2% of pregnant women attending ANC at Ain Shams University Hospital knew about the role of folic acid supplementation in prevention of congenital anomalies(16).A community based study done in Ethiopia had revealed that 27.5% of reproductive age women had good level of knowledge on preconception care(17).

Studies suggested antenatal care ought to initiate before pregnancy for a better pregnancy outcome. Implementation of preconception care in maternity care unit is crucial to achieve the sustainable development goal (SDG) targets in relation to maternal, neonatal and child health, by decision makers and stakeholders. However, evidence on the levels of knowledge and attitude on preconception care amongst women in rural African settings is scarce. The purpose of the study was therefore to measure the levels of knowledge and attitude on preconception care and their determinants among women who delivered at government hospitals in a rural setting in southern Ethiopia.

## Methods

### Study design and setting

A hospital-based cross sectional study was done from January 01to February 30, 2017 among mothers who delivered in public hospitals in Wolayita Zone and who were on immediate postnatal ward. Wolayita zone is found in the Southern Nations, Nationalities and Peoples Regional State of Ethiopia. According to the 2007 census of Ethiopia, the total population of the zone was 1.7 million. The public health institutions found in the zone were one referral hospital, four district hospitals and 70 health centers (5 urban and 65 rural). The total number of births from the five hospitals in 2016 was 7445 (Otona Hospital 3511, Bonbe hospital 1228, Halale hospital 1142, Bitana Hospital 956, and Bale Hospital 608).

### Study population and sampling procedures

Study populations were women who delivered at government hospitals in Wolayita zone during the study period. Mothers who were loss of consciousness, had mental problem, and referred to other hospitals were excluded.

Sample size was determined by using the software Epi Info version 7 with following assumptions: 95% confidence interval, an anticipated proportion of knowledge of preconception care of 10.4% based on a study in Nigeria(18), 4% of margin of error and a design effect of 1.5. The calculated sample size was 336. Added of 10% non-response rate, total sample was 374.

All public hospitals in Wolayita zone were included in the study and the sample size was proportionally allocated into five public hospitals based on number of deliveries each hospital. Systematic random sampling procedure was used to select study participants in each hospital. Monthly expected number of deliveries at public hospitals in Wolayita zone was 620; thus the sampling interval used was 2.

The questionnaires were prepared by reviewing the existing literatures. The questionnaire was prepared at English and then translated to Wolaytigna, and back to English to check uniformity. The questionnaire consisted of 57 items: 13 sociodemographic items, 6 obstetric items, 4 source of information items, 23 knowledge variables, and 11 attitude items. Attitude items level into five Likert scale (1-strongly disagree, 2-disagree, 3-neutral, 4-agree and 5-strongly agree). During analysis, the Likert scale items were categorized into three response categories to compute women’s attitude on preconception care: disagree (by merging 1-strongly disagree and 2-disagree), neutral and agree (by merging 4-agree and 5-strongly agree).

In Hawassa University Comprehensive Specialized Hospital pretest carried out 5% of study participant. Based on the pretest findings, amendment was done before initiation of actual data collection.

Data were collected using structured and pretested interviewer administered questionnaire through face -to -face by 10 midwives who had received training on basic emergency obstetrics and newborn care (BEmONC) and who can fluently communicate with the local language (Wolaytigna).Training was given to data collectors for three days on data collection methodology and related issues prior to the start of data collection time and were closely supervised during the data collection period.

### Statistical analysis

Data entry was done EPI Data 3.1 and transferred to SPSS version 20.0 for analysis. Based on 23 knowledge items, we computed an overall knowledge score for each study participant. Those who had knowledge score above the mean knowledge score were level as “adequate knowledge” whereas at or below the mean knowledge score were categorized as “inadequate knowledge”. Eleven attitude items were recorded into disagree, neutral and agree. Those whose response was “agree” were considered as having “positive attitude” towards preconception care, whereas those whose response was “disagree” were regarded as having “negative attitude” towards preconception care; those with a “neutral” response were considered as having “neither negative nor positive attitude”. Descriptive analysis was done to calculate and describe the basic characteristics of the study participants knowledge and attitude to preconception care. Binary logistic regression was used to identify the correlates of knowledge on preconception care, while ordinal regression was used to identify correlates of attitude towards preconception care. Adjusted odds ratios (AORs) with 95% confidence intervals (CIs) were used to judge the presence and strength of association between dependent and independent variables. P value <0.05 was taken as statistically significant.

## Results

### Socio-demographic characteristic of study participants

Three hundred seventy women participated in this study and 99% of response rate. The participants’ age were amid from fifty to thirty eight with a mean age of 25 (±4) years. Wolayita was the dominant ethnic group (91.9%). Three hundred sixty three (98.1 %) were married. The majority (69.7%) of the participants were housewives and 34.9 % had completed primary school (Table 1).

**Table 2:**
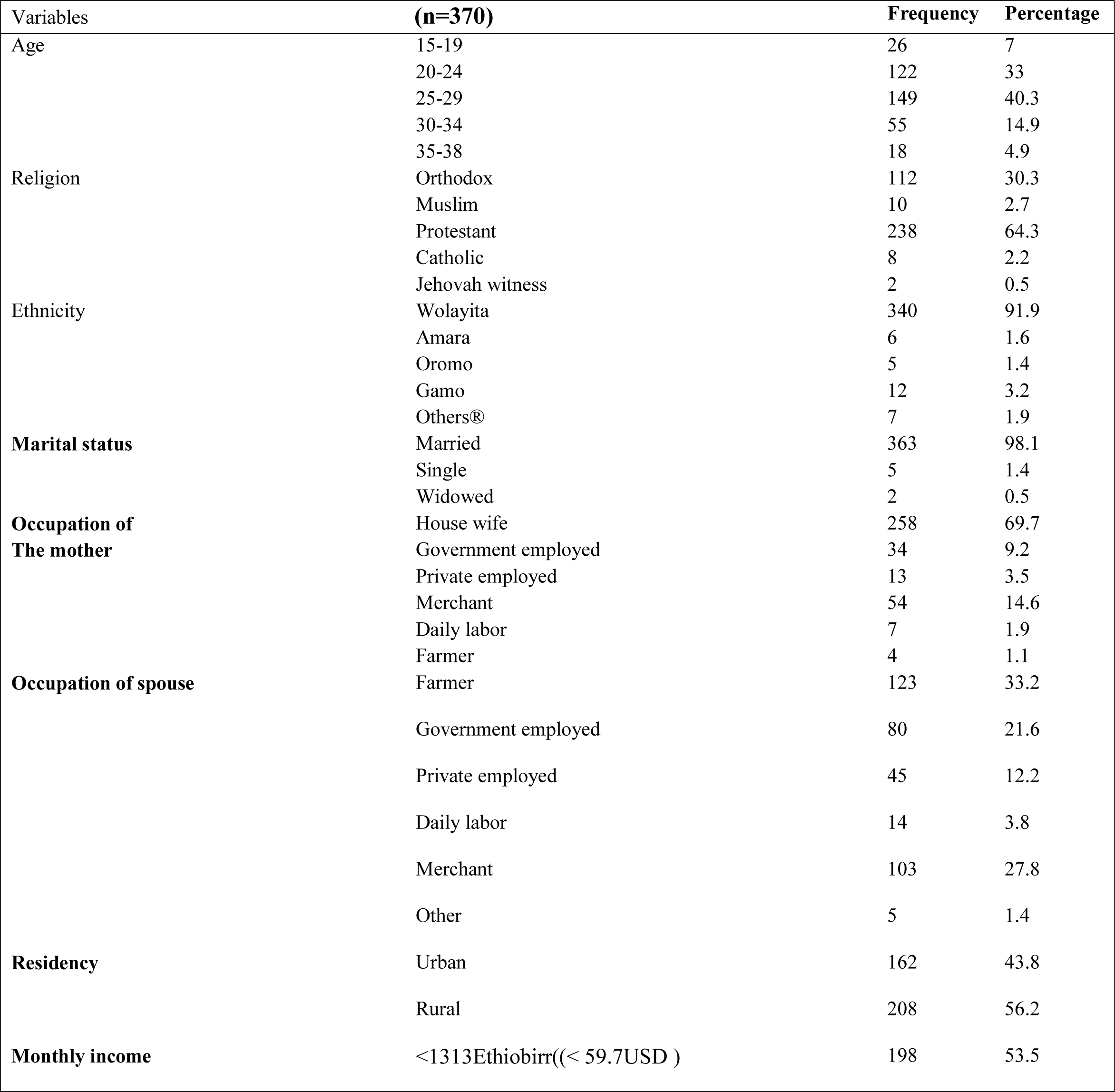
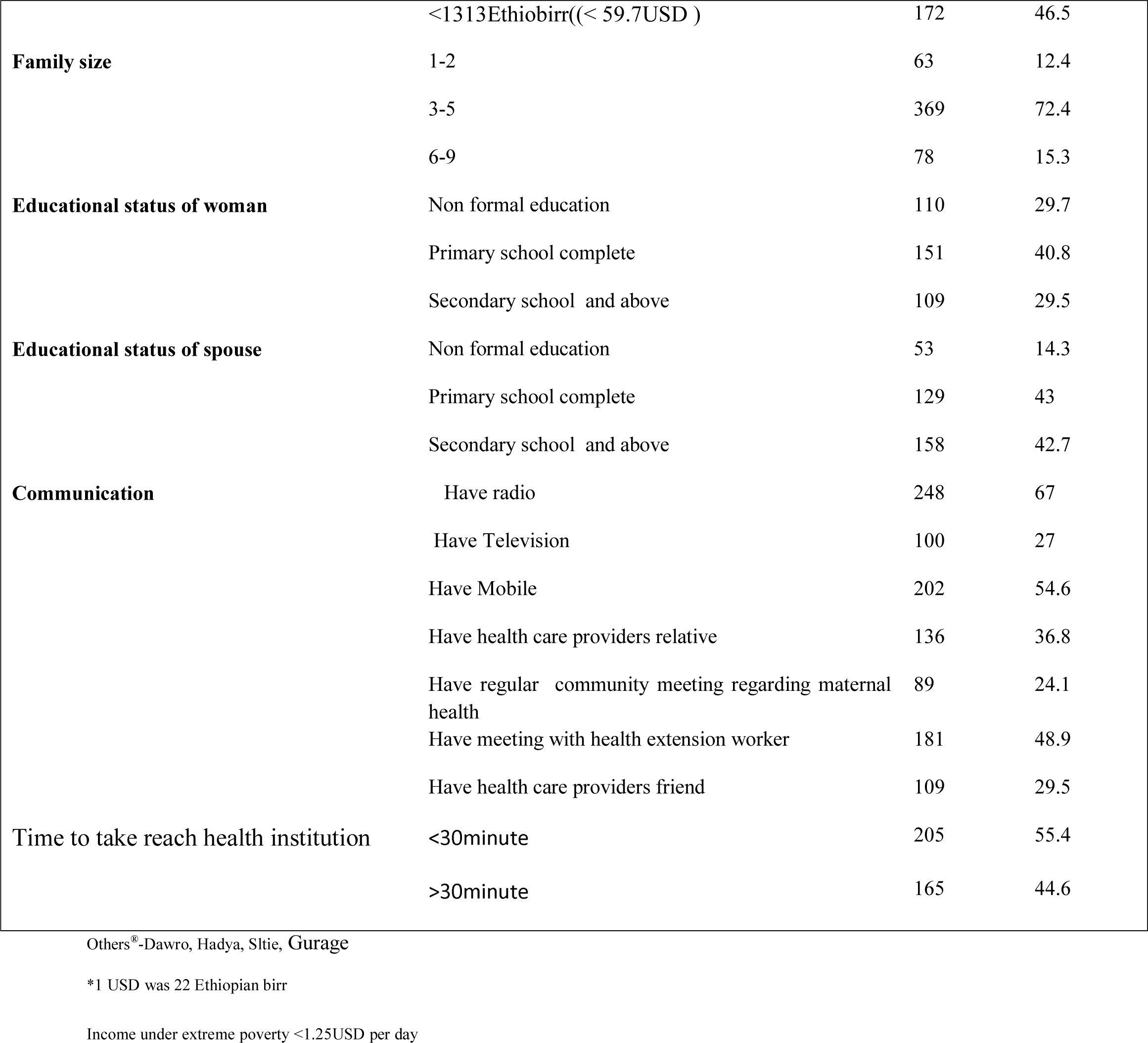
Socio-demographic characteristics of women who gave birth at govrmement hospitals in Wolayta Zone, South Ethiopia, February 2017

### Obstetric characteristics of study participants

In two hundred ninety six (80%) of the mothers, the recent pregnancy was planned. Nearly two-third (65.1%) of mothers had used family planning before the current pregnancy. Ninety eight (26.5%) of the mothers were primigravidae and 272(73.5%) were multigravidae, whereas 110(29.7%) were primipara and 260(70.3%) were multipara. Two hundred eighty three (76.5%) of the participants had antenatal contact for this pregnancy, of whom 152(41.1%) had four or more ANC contacts (Table3).

**Table 4:** Obstetric history of women who delivered at government hospitals in Wolayita Zone, South Ethiopia, February 2017

| Variables | (N=370) | Frequency | Percentage (%) |
| --- | --- | --- | --- |
| Have family planning use history | Yes | 241 | 65.1 |
|  | No | 129 | 34.9 |
| Gravida | Prim gravida | 98 | 26.5 |
|  | Multigravida | 272 | 73.5 |
| Parity | Primipara | 110 | 29.7 |
|  | Multipara | 260 | 70.3 |
| Is pregnancy plan | Yes | 296 | 80 |
|  | No | 74 | 20 |
| ANC follow up | Yes | 283 | 76.5 |
|  | No | 87 | 23.5 |
| Number of ANC visit | No visit | 20 | 5.4 |
|  | 1 | 9 | 2.4 |
|  | 2 | 44 | 11.9 |
|  | 3 | 130 | 35.1 |
|  | 4 | 152 | 41.1 |
|  | More than four | 15 | 4.1 |

### Level of mothers’ knowledge on preconception care

The lowest and highest knowledge scores of the mothers were zero to twenty three. One hundred ninety six (53%) (95% CI: 47.8%, 58.1%) of women had adequate level of knowledge to preconception care (Table 5). The main source of information were health institutions (33%) and friends (26.5%) (Figure 1).

**Table 6:** Women’s knowledge on preconception care who delivered at government hospitals in Wolayita Zone, South Ethiopia, February 2017

| Variable | ( N= 370) | Frequency | Percent |
| --- | --- | --- | --- |
| Avoid bad habits when planned to pregnancy | Yes | 311 | 84.1 |
|  | No | 59 | 15.9 |
| Adjust their life when planned to pregnancy | Yes | 324 | 87.6 |
|  | No | 46 | 12.4 |
| Avoid smoking when planned to pregnancy | Yes | 281 | 75.9 |
|  | No | 89 | 24.1 |
| Avoid drinking alcohol when planned to pregnancy | Yes | 291 | 78.6 |
|  | No | 79 | 21.4 |
| Avoid multiple sexual partners when planned to pregnancy | Yes | 303 | 81.9 |
|  | No | 67 | 18.1 |
| Test HIV/AIDS when planned to pregnancy | Yes | 302 | 81.6 |
|  | No | 68 | 18.4 |
| Take folic acid and multivitamins to prevent neural tube defects | Yes | 210 | 56.8 |
|  | No | 160 | 43.2 |
| Take iron sulfate to prevent anemia? | Yes | 293 | 79.2 |
|  | No | 77 | 20.8 |
| Avoid illicit drugs when planned to pregnancy | Yes | 262 | 70.8 |
|  | No | 108 | 29.2 |
| Stop over exercising when planned to pregnancy | Yes | 287 | 77.6 |
|  | No | 83 | 22.4 |
| Stop caffeine drinking when planned to pregnancy | Yes | 110 | 29.7 |
|  | No | 260 | 70.3 |
| Stop mercury from consumption of seafood when planned to pregnancy | Yes | 99 | 25.9 |
|  | No | 274 | 74.1 |
| Away from Pesticides/insecticides chemicals when planned to pregnancy | Yes | 217 | 58.6 |
|  | No | 153 | 41.4 |
| Away from contact with substances like lead in paints when planned to pregnancy | Yes | 102 | 27.6 |
|  | No | 268 | 72.4 |
| Away from exposure to occupational hazards when planned to pregnancy | Yes | 281 | 75.9 |
|  | No | 89 | 24.1 |
| Maintain body weight when planned to pregnancy | Yes | 241 | 65.1 |
|  | No | 129 | 34.9 |
| Take balance diet when planned to pregnancy | Yes | 266 | 71.9 |
|  | No | 104 | 28.1 |
| Check STI when planned to pregnancy | Yes | 301 | 81.4 |
|  | No | 69 | 18.6 |
| Take ordinary multivitamins when planned to pregnancy | Yes | 257 | 69.5 |
|  | No | 113 | 30.5 |
| Take ordinary vitamin D when planned to pregnancy | Yes | 112 | 30.3 |
|  | No | 258 | 69.7 |
| Take omega 3 vitamins when planned to pregnancy | Yes | 18 | 4.9 |
|  | No | 352 | 95.1 |
| Take ordinary zinc when planned to pregnancy | Yes | 18 | 4.9 |
|  | No | 352 | 95.1 |
| Street drugs when planned to pregnancy | Yes | 242 | 65.4 |
|  | No | 128 | 34.6 |

**Figure 2:**
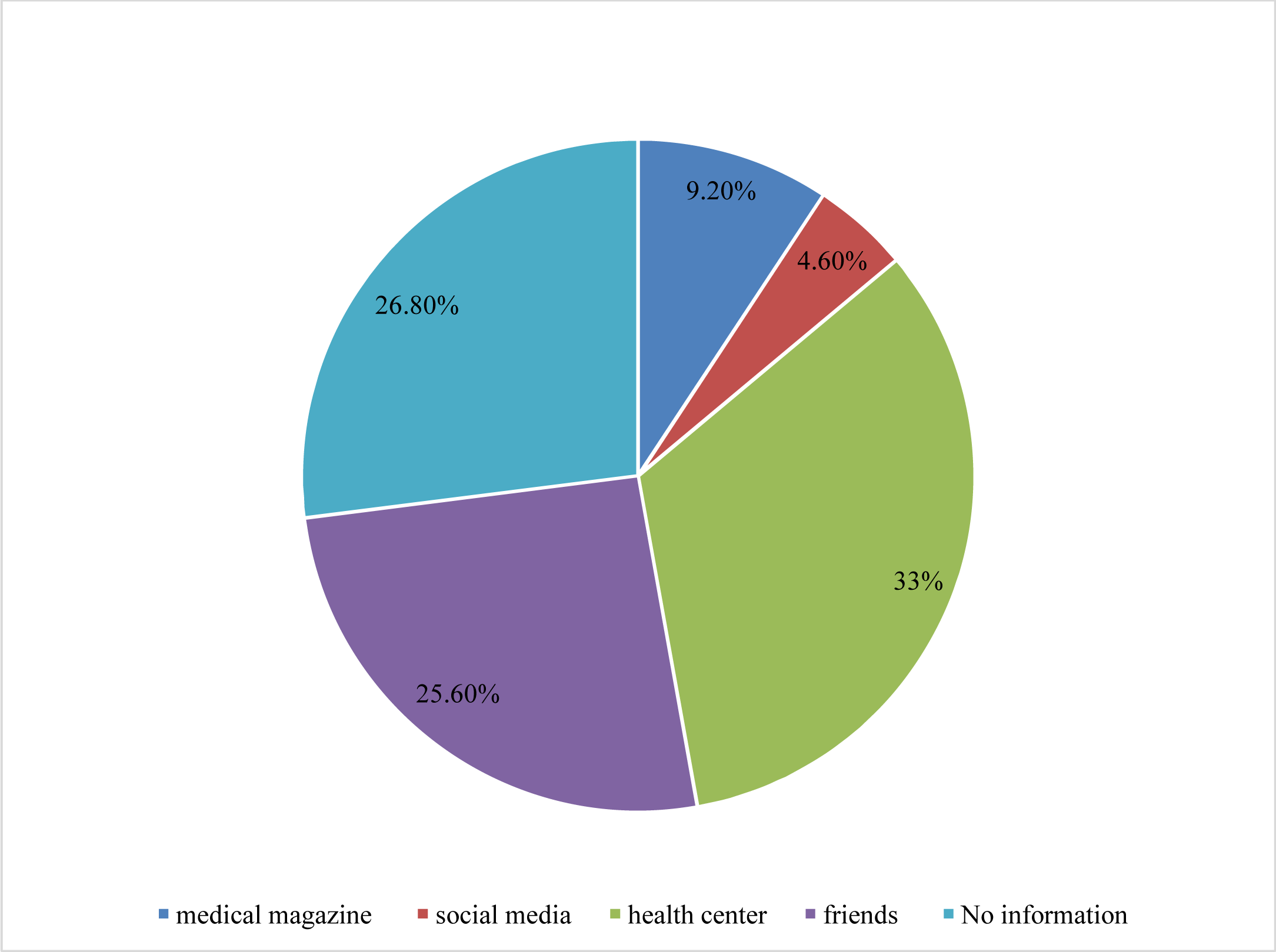
Source of information regarding preconception care amongst women who delivered at government hospitals in Wolayita Zone, South Ethiopia, February 2017.

### Women’s attitude on preconception care

Among the total of 370 respondents, 300(81.1%) of the mothers agreed that a hospital setting is the best place to provide preconception care and 277(74.9%) of women also agreed that preconception care is an important health issue for child bearing age women. Besides, 54(14.6%) of women agreed that there is not enough time to plan to get a preconception care. Overall, 201 (54.3%) (95%CI: 49.2%, 59.5%) of mothers had positive attitude towards preconception care, 23(6.2%) (95%CI: 4.1%, 8.9%) of mothers had neither positive nor negative (neutral) attitude towards preconception care and 146 (39.5%) (95%CI: 34.6%, 44.6%) of mothers had negative attitude towards preconception care (Table 7).

**Table 8:** Women’s attitude on preconception care who delivered at government hospitals in Wolayita Zone, South Ethiopia, February 2017

| Parameter (N= 370) | SA&A |  | Neutral |  | SD&D |  |
| --- | --- | --- | --- | --- | --- | --- |
|  | N | % | N | % | N | % |
| Preconception care does not have any effect on pregnancy outcome | 160 | 43.2 | 53 | 14.3 | 157 | 42.4 |
| Preconception care is an important health issue for women of child bearing age | 277 | 74.9 | 57 | 15.4 | 36 | 9.7 |
| A dedicated clinic for preconception care is a luxury service | 209 | 56.5 | 56 | 15.1 | 105 | 28.4 |
| A hospital setting is the best place to provide preconception care | 300 | 81.1 | 33 | 8.9 | 37 | 10 |
| Preconception care is a high priority all mother to plan pregnancy | 241 | 65.1 | 74 | 20 | 55 | 14.9 |
| I am not the most suitable person plan to get preconception care | 69 | 18.6 | 40 | 10.8 | 261 | 70.5 |
| There is not enough time to plan to get a preconception care | 54 | 14.6 | 43 | 11.6 | 273 | 73.8 |
| Health institutions exercise preconception care | 96 | 25.9 | 41 | 11.1 | 233 | 63 |
| Do you think high-risk mothers only start preconception care when planned to pregnancy | 106 | 28.6 | 28 | 7.6 | 236 | 63.8 |
| History congenital anomalies only use preconception care | 113 | 30.5 | 33 | 8.9 | 224 | 60.5 |
| Preconception care depends on health care providers willingness | 262 | 70.8 | 46 | 12.4 | 62 | 16.8 |

SA: Strongly agree, A: agree, SD: strongly disagree and D: disagree

### Determinants of knowledge and attitude on preconception care

Study participants who had radio (AOR: 2.91; 95% CI: 1.69, 5.43), planned pregnancy counterpart (AOR: 5.76; 95% CI: 2.84, 11.67), and had participated in community meetings related to preconception care (AOR: 2.96; 95% CI: 1.62, 5.43) had significantly higher odds of good level of knowledge on preconception care (Table 9).

**Table 10:** Determinants of knowledge on preconception care amongst women who delivered at government hospitals in Wolayita Zone, South Ethiopia, February 2017

| Variable | knowledgeable(N=196) | Non knowledgeable (N=174) | COR 95 % CI | AOR 95 % CI |
| --- | --- | --- | --- | --- |
| Do you have radio |  |  |  |  |
| Yes | 159 (43) | 89(24.1) | 4.10(2.58,6.54) * | 2.91(1.69,5.43) * |
| No | 37(10) | 85(23.4) | 1 | 1 |
| Do have health care providers' relatives |  |  |  |  |
| Yes | 88(23.8) | 48(13) | 2.13(1.384,3.306)* | 1.29(0.74,2.26) |
| No | 108(29.2) | 126(34.1) | 1 |  |
| Is pregnancy planned |  |  |  |  |
| Yes | 183(49.5) | 113(30.5) | 7.60(3.995,14.455)* | 5.76(2.84, 11.67) * |
| No | 13(3.5) | 61(16.5) | 1 | 1 |
| do you have community meeting related to preconception care |  |  |  |  |
| Yes | 67(18.1) | 22(5.9) | 3.588(2.100,6.132) * | 2.96(1.62, 5.43) * |
| No | 129(34.9) | 152(41.1) | 1 | 1 |
| Do you have health care |  |  |  |  |
| providers friends |  |  |  |  |
| Yes | 75(20.3) | 34(9.2) | 2.552(1.591,4.094) * | 1.36(0.74,2.47) |
| No | 127(34.3) | 140(37.8) | 1 |  |
| Educational status of spouse |  |  |  |  |
| Non formal education | 14(3.8) | 39(10.5) | 0.301(0.151,0.597) |  |
| Primary school | 96(25.9) | 63(17) | 1.28(0.817,1.993) * | 1.31(0.73,2.36) |
| Secondary and above | 86(23.2) | 72(19.5) | 1 | 1 |

\* P<0.05

On other hand, multivariable ordinal regression showed that women who had mobile phone had 2 times higher chances of positive attitude(AOR: 2.17, 95 % CI: 1.31, 3.59) and those who had participated in community meetings related to preconception care had decreased odds of positive attitude towards preconception care (AOR: 0.36, 95 % CI: 0.22, 0.60) (table 11).

**Table 12:**
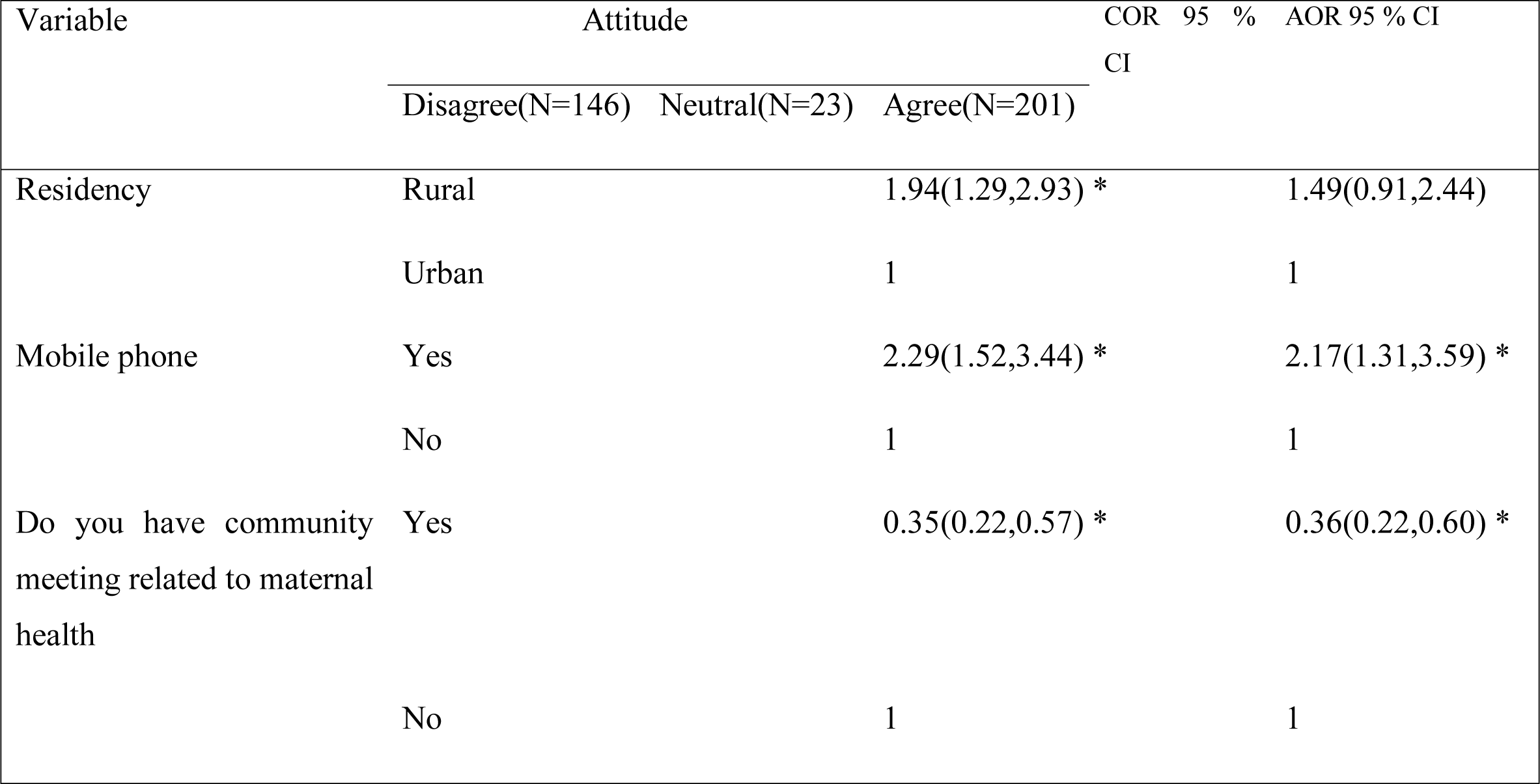
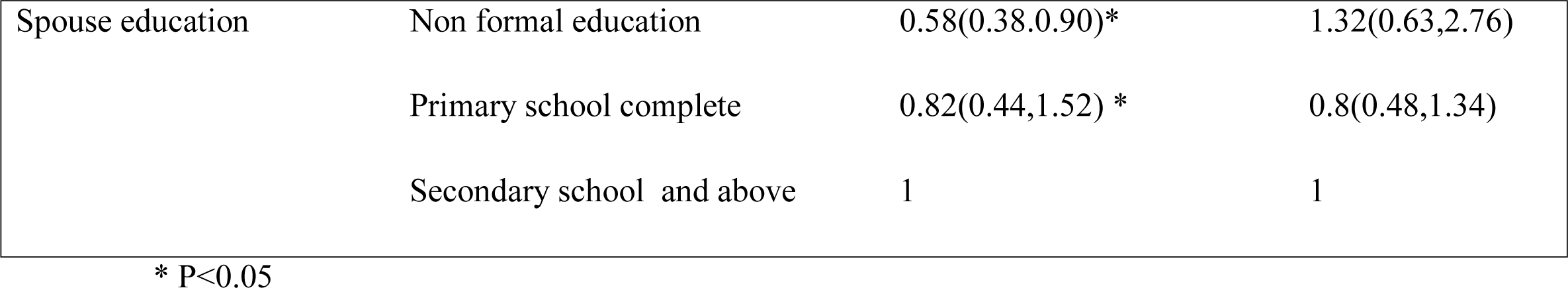
Determinants of attitude to preconception care amongst women who delivered at government hospitals in Wolayita Zone, South Ethiopia, February 2017

## Discussion

Findings revealed that level of knowledge on preconception care amongst women who delivered at government hospitals in Wolayita Zone is 53%. This finding is inconsistent with the findings in Northwest Ethiopia (27.5%) (17), Sudan (11.1%) (19), Nigeria (2.5%)(9), Iran (10.4%)(20), Saudi Arabia (37.9%) (21), United Arab Emirates (46.4%) (22), and Turkey (46.3%) (23). The possible explanation for higher level of knowledge in the present study could be the time of study currently, maternal health is given high attention which may result in an overall increase in knowledge of issues related to maternal health. Contextual differences in the study settings could also account for the observed differences.

On the other hand, it is consistent with studies done in Malaysia (51.9%)(15), and in Qatar (53.7 %)(24). However, this fining is lower than the study done in Canada (70%)(25), Jordan (85%)(26), British Colombia (71%) (27), Saudi Arabia (84.6%)(28), and in the United States of America (76%) (29). The possible explanation could be low level of knowledge due to health sector infrastructure difference, socioeconomic difference, lack of health wellness clinic in the area of the present study, lack of preconception service across Ethiopia, lack of promotion on preconception care by mass media, and low commitment of health care providers due to high load of clients.

In this study the correlates of knowledge on preconception care were found to be possession of transistor radio, planned pregnancy, and having participated in community meetings related to preconception care. Women who had radio had three times higher likelihoods to have adequate knowledge on preconception care. It is inconsistent with studies done in Ethiopia and Nigeria(9, 17). The higher level of knowledge of preconception care amongst women who possess transistor radio and who participate in community meetings related to preconception care can be due to exposure of such mothers for health information via radio and also during community meetings. The community meetings could also create a platform for women to share their positive and negative childbirth experience and prevention mechanisms. Similarly, women who planned the recent pregnancy had six times higher chances to have adequate knowledge on preconception care which is coincided with the finding in Brazil(30). The possible explanation could be reproductive age women who planned pregnancy are anticipated to know their healthiness correlated to maternal health care and may thus have also a better awareness of issues correlated to preconception care.

In this study 54.3% of mothers were found to have positive attitude on preconception care. This finding is incomparable with studies done Malaysia (98.5%)(15)and USA (98%) (29).The difference might be due availability and accessibility of the service in settings with better socioeconomic status such as in Kelantan, Malaysia and USA.

Women who possess mobile cell phones have more than twice increased odds of positive attitude towards preconception care; however, women who have participated in community meetings related to preconception care had decreased odds of positive attitude towards preconception care. The reason why women who possess cell phone have a higher odds of positive attitude towards perception care could be due to better exposure of such women to health information via frequency modulated (FM) radio services which are available in most cell phones and for some of the literate mothers via mobile internet. Women who posses mobile phone may also generally be in a better socioeconomic position and hence may have more positive attitudes to health care services. Why women who participate in community meetings have a decreased odds of positive attitude is difficult to explain, but could be a result of being fed up with regular participation in community meetings.

The strength of this study relative to previous studies is incorporating relevant variables which were not addressed previously such as having planned pregnancy, possession of transistor radio and participating in community meetings related to preconception care. The limitation of this study is point out that it did not incorporate both sides like partners of women. Outcomes can be some amount affected by recall and social desirability biases.

## Conclusion

Levels of women’s knowledge and positive attitude on preconception care among women who delivered at government hospitals in rural southern Ethiopia is low compared with other studies. Using transistor radio and mobile phone have significant effects in improving the knowledge and attitude of reproductive age women on preconception care. Hence, providing community health education based on radio and/or mobile phone messaging could be useful in positively influencing the knowledge and attitude of women on preconception care.

## Declarations

## Abbreviation

AOR: Adjusted Odds Ratio
BEmONC: basic emergency obstetrics and newborn care
CI: confidence interval
SDG: Sustainable Development Goals
SPSS: Statistical Package for Social Sciences
USA: United States of America

## Acknowledgement

We would like to acknowledge the Centre of Excellence in Maternal and Newborn Health SRH3 project at the College of Medicine and Health Sciences of Hawassa University for technical and financial support. We extend to acknowledge study participants, Wolayta Zone Health office and Wolayta Zone public hospital officials for their support and permission to carry out the study.

## Authors’ contribution

ZY originated of the idea and planned to the study, participating during data collection, analysis the data and write up the manuscript. ZT, AA, MS, GB, KT, SM and ZK are reviewed the study procedure, participated in data acquisition and analysis and reviewed the manuscript. All authors read and approved the final manuscript.

## Ethics and consent

Ethical clearance was gained from the Institutional Review Board at the College of Medicine and Health Sciences of Hawassa University. Wolayita Zone Health office and management of the respective public hospitals gave consent to conduct the study. Written consent was obtained from the study participants before data collection started. Anonymous questionnaire was used to assure confidentiality of study participants.

## Fund

This study sponsored by Hawassa University, Centre of Excellence in Maternal and Newborn Health. The funders had no role in the study design, data collection, data analysis, data interpretation, or writing of the report. The authors had access to the data in the study and the final responsibility to submit the paper.

### Consent for publication

Not applicable.

## Availability of data and materials

All data on which this article is based are included within the article.

## Authors’ information

1. **Zemenu Yohannes:** School of Nursing and Midwifery, College of Medicine and health sciences, Hawassa University, Ethiopia
2. **Zelalem Tenaw:** School of Nursing and Midwifery, College of Medicine and health sciences, Hawassa University, Ethiopia
3. **Ayalew Astatkie:** School of Public Health, College of Medicine and health sciences, Hawassa University, Ethiopia
4. **Melesse Seyoum:** School of Nursing and Midwifery, College of Medicine and health sciences, Hawassa University, Ethiopia
5. **Gezahegn Bekele:** School of Nursing and Midwifery, College of Medicine and health sciences, Hawassa University, Ethiopia
6. **Kefyalew Taye:** School of Medicine, College of Medicine and health sciences, Hawassa University, Ethiopia
7. **Sewangizaw Mekonnen:** School of Nursing and Midwifery, College of Medicine and health sciences, Hawassa University, Ethiopia
8. **Zerai Kahsaye:** School of Medicine, College of Medicine and health sciences, Hawassa University, Ethiopia

